# Hypothetical control of postural sway

**DOI:** 10.1101/2020.05.19.104760

**Authors:** Madhur Mangalam, Damian G. Kelty-Stephen

## Abstract

Quiet standing exhibits strongly intermittent variability that has inspired at least two interpretations. First, variability can be intermittent through the alternating engagement and disengagement of complementary control processes at distinct scales. A second and perhaps deeper way to interpret this intermittency is through the possibility that postural control depends on cascade-like interactions across many timescales at once, suggesting specific non-Gaussian distributional properties at different timescales. Multiscale probability density function (PDF) analysis shows that quiet standing on a stable surface exhibits a crossover from low, increasing non-Gaussianity (consistent with exponential distributions) at shorter timescales, reflecting inertial control, towards higher non-Gaussianity. Feedback-based control at medium to longer timescales yields a linear decrease characteristic of cascade dynamics. Destabilizing quiet standing with unstable surface or closed eyes serves to attenuate inertial control and to elicit more of the feedback-based control over progressively shorter timescales. The result was to strengthen the appearance of the linear decay indicating cascade dynamics. Finally, both linear and nonlinear indices of postural sway also govern the relative strength of crossover or of linear decay, suggesting that tempering of non-Gaussianity across log-timescale is a function of both extrinsic constraints and endogenous postural control. These results provide new evidence that cascading interactions across longer-timescales supporting postural corrections can even recruit shorter-timescale processes with novel task constraints that can destabilize posture.

## 1. Introduction

### 1.1. Postural sway in quiet standing

Quiet standing—a fundamental motor act supporting other movement tasks—exhibits intermittent variability. This intermittency can be interpreted in various ways. First, variability can be intermittent due to the engagement and disengagement of control processes. For instance, during quiet standing, the body continually sways about a constant average position of the global ground reaction force (center of pressure, CoP) trajectory. Within the center of the base of upright support, it is traditionally thought that the postural system relies on postural corrections based on sensory feedback at longer timescales, towards the margins of the base of support [1,2]. A second and perhaps a deeper way to interpret this intermittency is through the possibility that feedback-based corrections depend on the cascade-like interactions across timescales. For instance, more modern interpretations of postural control recognize that upright standing is not always about reining the body within a margin. Instead, postural control appears to respect the maintenance of equilibrium points around which the body can ‘tremble,’ but postural control can also ‘ramble,’ sometimes retaining equilibrium points in place but sometimes moving those about [3,4]. Quiet standing accomplishes its rambling equilibrium through the integration of information from visual, vestibular, and somatosensory receptors, in conjunction with the passive properties of the musculoskeletal system. The neurological and physiological processes that have been suggested to dictate intermittency in postural sway include active mechanical processes [5–7], passive reflex mechanisms [8,9] anticipatory [10–12] and feedback processes [13,14], and exploratory processes used by the central nervous system (CNS) to generate sensory information [15,16].

The sources of intermittency noted above should each entail characteristic signatures on the distributional properties of postural sway. Different modes of postural control enlist different kinds of relationships between consecutive fluctuations. For instance, feedback-based correction will entail a temporal structure that feedforward fluctuations may not, depending on how much independence postural control might leave between fluctuations. The change in expected independence will entail a change in expected distributions. An important theme in what follows is aligning the transition in control with a transition in distribution of measured postural fluctuations.

### 1.2. Aligning feedback-based control with multiplicative random processes and non-Gaussianity

The transition from more to less independence among postural fluctuations may prompt the transition from more Gaussian to less Gaussian (i.e., to more non-Gaussian) probability density functions (PDFs). As independence gives ways to interactivity, the statistical processes that these postural fluctuations resemble move from additive to multiplicative. Independence aligns with additivity, and interactivity aligns with multiplicative processes. First, postural sway remaining within the base of support is most compatible with independence among postural fluctuations. Under these conditions, feedback-based correction becomes a lower priority for maintaining an upright stance, especially as a precision suprapostural task begins to take higher priority [17]. Specifically, shorter-timescale sway roams ‘unchecked’ resembling Brownian motion [1,2], exhibiting inertia-driven temporal correlations across fluctuations whose nonzero size is drawn from distributions with thin tails (e.g., exponential) [18,19]. The independence of postural fluctuations from feedback in this inertial mode ensures an additive random process for which Gaussian variance effectively describes sway. Nearer to the boundaries of the support surface, the postural system draws on feedback to maintain stability, correcting for and so reversing the inertial tendencies. Feedback is fundamentally contingent and interactive, leading postural fluctuations to exhibit lognormal distributions characteristic of multiplicative random processes [20,21]. The lognormal PDFs’ heavy tails entail that a non-Gaussian variance better describe sway. That is, stable quiet standing should exhibit exponential and lognormal distributions at shorter and longer timescales devoted to inertial and correction-based control modes, respectively. Variance requires different definitions for lognormal and Gaussian distributions, and the non-Gaussianity of multiplicative processes manifests as an excess of lognormal variance beyond Gaussian variance.

An important step beyond the simple additive-vs.-multiplicative dichotomy is our proposal that feedback-based corrections are explicitly cascade-like—multiplicative random processes with hierarchical interactivity [22–24]. Feedback is necessarily a consequence of outward movements of the organism into the surrounding task context, and hence it is always sequential—organism activity always prompts later feedback. The question of the hierarchical structure only enters into this sequence as a potential nuance.

The sparsest series of feedback may show no cascade structure. That is, feedback could straightforwardly be a chain of sensorimotor links (sensory events only following motor events, and vice versa). Time-series modeling often refers to sequences with feedback only on previous events as having ‘short memory’ [25]. The popularity and controversy around heavy-tailed distributions warrants emphasis that this extremely short memory is a valid interpretation of interactivity from non-Gaussian distributions [26–29]. Many heavy-tailed distributions result from little more than outliers and so have not even the least meaningful sequential relationship [30]. Lognormal and other heavy-tailed distributions are a signature of what could be feedback and more than short-memory feedback, but we cannot take these interpretations for granted without further test. We can conceive of longer-memory feedback following from longer links in this chain, e.g., a brief sensory event (e.g., an action potential) that could follow short and long motor events (e.g., a muscle fiber contracting and the extension of a joint, respectively). Longer-term, less-motor, and more-cognitive factors like intention or instructions might contribute as well to the sensory event. So, we can conceive at least three lengths of links: the shortest muscle-fiber link, the slightly longer joint-extension link, and the longest intention link. Feedback can follow from a mix of long-memory and short-memory without entailing cascade organization [23,24].

### 1.3. Empirical test of cascades in feedback-based postural control

Multiplicative processes are cascade-like when longer events can reshape shorter events. It would be wrong to think of cascades and short-memory as exclusive opposites; cascades can exhibit short-memory chain structure [31,32]. What distinguishes cascades is a hierarchical structure affording not just short-memory but extensions including long-memory and more fluid, flexible organizations of feedback.

The cascades reflect multiplicative random processes in which the memories at different timescales intermix. They allow mathematically addressing situations in which feedback is deeply context-sensitive. For instance, during quiet stance, the value of feedback may change depending upon how intention, joint extension, and muscle-fiber activity influence each other. In these situations, the cascade formalism reflects an accrual of possible feedback regimes, not sloughing the other simpler kinds. So, the preceding examples of feedback are available as limiting cases of more generic cascade mechanisms [32]. The ‘chain’ metaphor above eventually becomes too rigid to describe cascades respecting the fact that dynamics have been used to model fluid dynamics (e.g., actual, hydrological cascades [33]). The interactions amongst links at different scales entail that nonlinearity is the rule than an exception to a linear sequence.

Nonlinear interactions across scales unleash a deep potential of cascades to invite a wider class of mathematical models of cause-effect relationships. As nonlinearity brings pliability to once-rigid links in our chain, cascades begin to look much less like chains and more like tapestries [34–36], branching structures [37– 41], nestings [42–44], or webs [45–47]. This metaphoric diversity mainly illustrates how nonlinear interactions across scales open up the scientific discourse to various proposals of how hierarchical geometry may (or may not) constrain the shape of cause-effect relationships. In the context of human postural control, cascade modeling allows testing whether preceding longer-memory links can all reshape one another as well as that brief sensory event called feedback. But then, we might note that feedback might be no less multi-layered, including an action potential in a sensory cell and longer-term perceptual impressions that the postural control system can use in the future.

Cascade-like multiplicative processes offer a generic, common analytical thread to coordinate this exploding range of possible forms for cascading feedback processes: ‘multifractal geometry.’ While cascades may fire the scientific imagination and allow us to tailor our modeling to different model systems, there is a promise that the widely diverse manifestations of nonlinear interactions across scales might operate along with similar multifractal principles [22,48,49]. The present work aims to develop the topic of human postural control through a multifractal lens to determine the sensitivity of cascade-formalisms to human postural control.

The present work applied multiscale PDF analysis rather than simply examining the ambiguous evidence of heavy tails. Multiscale PDF elaborates the multifractal formalism for quantifying the strength and form of cascades via quantifying non-Gaussianity. We tested for a crossover from Gaussianity (and exponential distribution) at shorter timescales to non-Gaussianity (and lognormal distribution) at longer timescales. To test whether the crossover was specific to the timescale-dependence of postural corrections, we examined multiscale non-Gaussianity under manipulations intended to destabilize posture. We hypothesized that destabilizing posture would elicit non-Gaussian postural corrections into the short timescales not traditionally associated with corrections in response to feedback. Multiscale PDF results indicate cascade dynamics when the full distribution of measured fluctuations exhibits a linear decay of non-Gaussianity with log-timescale. We compared original data to surrogate data to ensure that these cascade-like results depended on nonlinear interactions across scales [50–52].

### 1.4. Multiscale probability density function (PDF) analysis of postural sway in quiet standing

A central feature of intermittent, cascading fluctuations is the heterogeneity of variance beyond the bounds of a Gaussian probability distribution function (PDF). An important question then is about the expected PDF shape of a system whose intermittency reflects the cascading of fluctuation across many scales. Research on intermittency in the turbulent fluid flow [53] has inspired a method for quantifying this heterogeneity of variance through a metric 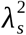 [54,55]. This metric speaks directly to a point raised above: multiplicative random processes will generate a lognormal PDF with larger variance than the Gaussian PDF resulting from additive random processes. As we had noted, variance calculated for Gaussian PDFs underestimates lognormal variance for multiplicative random processes. 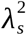 expresses the excess of lognormal variance compared to Gaussian variance. Hence, 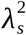 is zero only for Gaussian PDFs and increases as for progressively non-Gaussian PDFs.

Multiscale PDF analysis aims to distill deep insights about the random process producing measured behavior via empirical calculations of variance. Empirical variance can be unstable [56], depending on contingent aspects of measurement (e.g., sample size). Multiscale PDF analysis quantifies 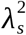 at a variety of timescales, coarse-graining measured fluctuations for progressively longer timescales. Controlling for how coarse-graining might depress variance, multiscale PDF reveals crucial features of cascades based on how quickly or slowly 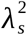 decays with timescale [57]. At shorter and medium scales, higher 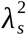 indicates stronger evidence of cascades. Theoretical simulations with idealized cascades show monotonic decay of 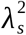 with longer timescales. Additionally, distinct kinds of cascades show distinct types of decay. For scale-invariant cascades, 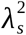decreases linearly with the natural logarithm of timescale (log-timescale), but cascades unfolding across a limited range of timescales exhibit quadratic decay with log-timescale [52].

This issue of scale-dependency motivates our attempt to manipulate the transition between inertial and feedback-based postural control by destabilizing the support surface of closing the eyes. This transition is scale-dependent—that is, it depends on the width of the base of support. We expected that the short-scale postural sway roaming inertially unchecked through the base of support would show low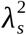, and higher 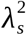only at medium timescales. Thus, we expected that the scale of the base of support would produce a quadratic shape, depressing the 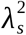-vs.-log-timescale curve over the short and long timescales. Finally, we expect that destabilizing the support surface—which effectively narrows the base of support and widens the range of timescales—or closing the eyes would reduce the scale-dependence of 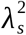 and resemble more the linear decay associated with scale-invariant cascades.

While the manipulations of support surface or vision might make the 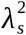-vs.-log-timescale curve less quadratic and more linear, their effects must depend on endogenous postural control. Multiscale PDF analysis of both heart-rate variability (HRV) [54,55,58–60] and neuronal interspike intervals (ISI) [61] suggests that endogenous control processes account for the variability in 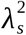-vs.-log-timescale curve for physiological dynamics. For instance, endogenous sinoatrial dynamics could explain the heavier-tailed PDFs for pathological cases than for healthy cases in HRV [62], and non-Gaussianity showed relationships with the magnitude autocorrelation of the ISI [61]. Hence, in addition to testing whether postural corrections at different timescales in quiet standing change the non-Gaussianity of CoP dynamics, we expected that endogenous variability in postural control would moderate any observed effect of these manipulations on the non-Gaussianity of postural fluctuations.

Here, we applied multiscale PDF method to study postural control and test three specific hypotheses in a reanalysis of a public dataset of kinematics and ground reaction forces [63]. Hypothesis-1 was that quiet standing on a stable surface would show a crossover between low non-Gaussianity in postural sway (consistent with exponential distributions) at shorter timescales reflecting lack of sensory corrections, increasing towards higher and subsequently decreasing (consistent with cascade-driven lognormal distributions) at longer timescales. Specifically, we tested for this crossover as a negative quadratic polynomial with log-timescale. Hypothesis-2 was that destabilizing quiet standing would require postural corrections at more of the shorter timescales, eliciting non-Gaussianity that begins high and decreases across timescales (consistent with cascade-driven lognormal distribution). Specifically, we expected that quiet standing on an unstable surface or with eyes closed would cancel the negative quadratic polynomial. Hypothesis-3 was that known indices of postural sway would significantly moderate the quadratic polynomial, effectively governing the appearance or disappearance of the crossover.

## 2. Methods

The present work is a reanalysis of a public data set with ground reaction forces of human balance [63].

### 2.1. Participants

Forty nine participants grouped by age: young (15 males and 12 females, 18–40 years old) and old (11 males and 11 females, > 60 years old) took part in the present study after providing institutionally-approved informed consent.

### 2.2. Experimental setup and procedure

Each participant stood still for 60 s under all four possible combinations of vision (eyes open/eyes closed) and standing surface (stable/unstable). Each participant completed three 60-s trials per condition. Stable condition involved standing on two 40×60 cm force platforms (100 Hz, OPT400600-1000; AMTI Inc., Watertown, MA) under each foot, and unstable condition involved standing on two 6-cm high foam blocks (Balance Pad, Airex AG Inc., Sins, Switzerland), each placed on each force platform.

### 2.3. Data processing

All data processing was performed in MATLAB 2019b (Matlab Inc., Natick, MA). Trial-by-trial ground reaction forces yielded a 2D center of pressure (CoP) series, each dimension describing CoP position along anterior-posterior (*AP*) and medial-lateral (*ML*) axes. Each 60-s trial yielded 6000-sample 2D CoP series and 5999-sample 2D CoP displacement series. Finally, a 1D CoP planar Euclidean displacement (PED) series described postural sway along the transverse plane of the body. Iterated Amplitude Adjusted Fourier Transformation (IAAFT) provided phase-randomized surrogates using original series’ spectral amplitudes to preserve only linear temporal correlations [23,64]. We used the IAAFT surrogate data for two purposes. Primarily, we computed multiscale PDF analysis on the IAAFT surrogates because any difference in the 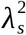-vs.-log-timescale curves between the original and IAAFT series would reveal differences in non-Gaussianity attributable to nonlinear temporal correlations. Secondarily, as noted below (Section 2.4), we wanted to test whether the difference in multifractal spectrum width due to nonlinear temporal correlations would moderate the non-Gaussianity in original series.

### 2.4. Canonical indices of postural control

Three linear indices were computed for each CoP PED series: (i) Mean of all fluctuations (CoP_ *Mean*), (ii) Standard deviation (CoP_*SD*), and (iii) Root mean square estimate (CoP_*RMSE*). These linear indices quantify the extent of and variation in postural sway with no reference to how the magnitude of sway at a given instance is related to the magnitude of away at other instances. Postural control under unstable conditions is associated with a higher magnitude of and variation in sway [65–67].

Four nonlinear indices were computed for each CoP PED series: (i) Sample entropy (CoP_ *SampEn*), indexing the extent of complexity in the signal [68]. We used *m* = 2 and *r* = 0.2 for computing the *SampEn* [cf. 34]. (ii) Multiscale sample entropy (CoP_*MSE*), indexing signal complexity over multiple timescales [70]. We used *m* = 2, *r* = 0.2, and *τ* = 20 for computing the *MSE* [cf. 34]. *SampEn* and *MSE* measure the repeatability or predictability (i.e., uncertainity) of the possible configurations in a time series. *MSE* extends *SampEn* to multiple timescales and, providing an additional perspective when the timescale of relevance to the behavior of interest is unknown. Structural changes in postural sway revealed by *SampEn* and *MSE* have lent insights into the effects of balance training, vision, and support surface on postural control [71,72]. (iii) We used detrended fluctuation analysis [73] to compute Hurst’s exponent, *H*_fGn_, indexing monofractal scaling or the strength of temporal correlations in original CoP PED series and shuffled versions of each series (CoP_*H*_fGn_) using the scaling region: 4, 8, 12,… 1024 [cf. 37]. Shuffling destroys the temporal structure of a signal. (iv) Multifractal spectrum width, Δ*α*, indexing multifractality or the extent of multifractal temporal correlations in the signal. Chhabra and Jensen’s direct method [74] was used to compute Δ*α* for the original CoP series and the IAAFT surrogate (CoP_dAlpha) [cf. 38]. Monofractal scaling and multifractal spectrum width have been traditionally implicated in quantifying the complexity in postural sway and its decay with aging and disease [76–78].

### 2.5. Multiscale probability density function (PDF) analysis

Multiscale PDF analysis characterizes the distribution of abrupt changes in CoP PED series {*b* (*t*)} using the PDF tail. The first step is to generate {*B* (*t*)} by integrating {*b* (*t*)} after centering by the mean *b*_*ave*_ (figure 1*a*):

**Figure 1.**
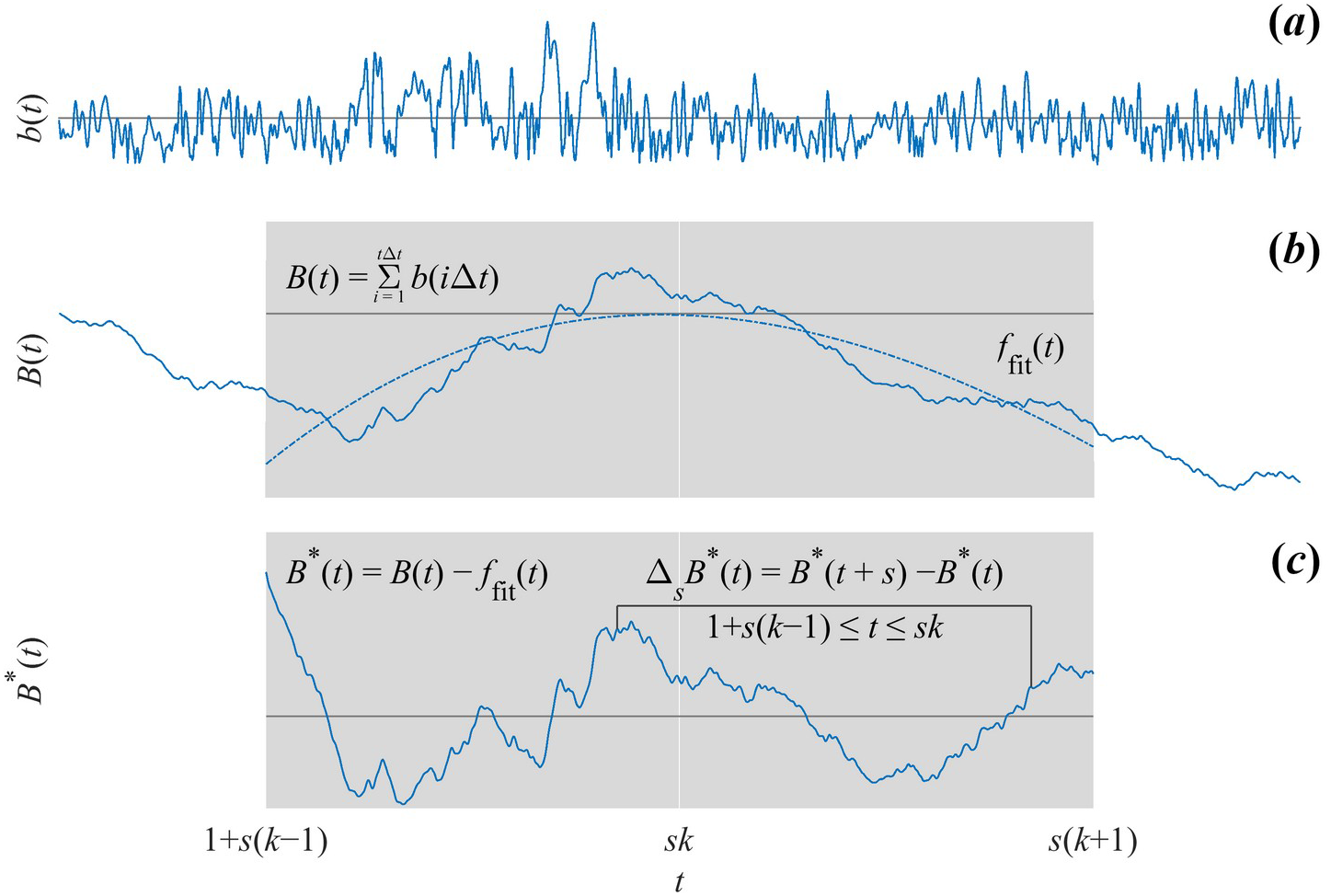
Schematic illustration of multiscale probability density function (PDF) analysis. (*a*) The first step is to generate {*B* (*t*)} by integrating {*b* (*t*)} after centering by the. 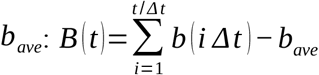 (*b*) A a 3^rd^ order polynomial detrends {*B* (*t*)} within *k* overlapping windows of length 2 *s, s* being the timescale. (*c*) Intermittent deviation *Δ*_*s*_ *B* (*t*) in *k* ^*th*^ window from 1+ *s* (*k −* 1) to *sk* in the detrended time series {*B*^*d*^ (*t*)=*B* (*t*) *− f* _*fit*_ (*t*)} is computed as *Δ*_*s*_ *B*^□^ (*t*)=*B*^□^ (*t* + *s*) *− B*^□^(*t*), where 1+ *s* (*k −* 1) ≤*t* ≤*sk* and *f* _*fit*_ (*t*) is the polynomial representing the local trend of {*B* (*t*)}, of which the elimination assures the zero-mean probability density function in the next step.

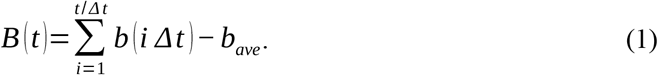

A 3^rd^ order polynomial detrends {*B* (*t*)} within *k* overlapping windows of length 2 *s, s* being the timescale (figure 1*b*). Intermittent deviation *Δ*_*s*_ *B* (*t*) in *k* ^*th*^ window from 1+ *s* (*k −* 1) to *sk* in the detrended time series {*B*^*d*^ (*t*)=*B* (*t*) *− f*_*fit*_ (*t*)} is computed as *Δ_s_ B*^*d*^ (*t*)=*B*^*d*^ (*t* +*s*) *− B*^*d*^ (*t*), where 1+ *s* (*k −* 1) ≤*t* ≤*sk* and *f*_*ave*_ (*t*) is the polynomial representing the local trend of {*B* (*t*)}, of which the elimination assures the zero-mean probability density function in the next step (figure 1*c*). Finally, *Δ*_*s*_ *B* is normalized by the SD (i.e., variance is set to one) to quantify the PDF.

To quantify the non-Gaussianity of *Δ*_*s*_ *B* at timescale *s*, the standardized PDF constructed from all the *Δ*_*s*_ *B* (*t*) values is approximated by the Castaing model [53], with *λ* _*s*_ as a single parameter characterizing the non-Gaussianity of the PDF. *λ*_*s*_is estimated as

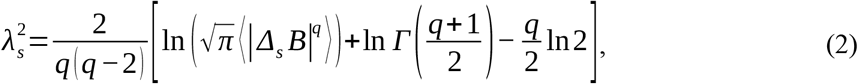

where ⟨|*Δ*_*s*_*B*|^*q*^ ⟩ denotes an estimated value of *q*^*th*^ order absolute moment of {*Δ*_*s*_; *B*}. As 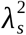 increases, the PDF becomes increasingly peaked and fat-tailed (figure 2*a*). 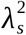 can be estimated by Eq. (2) based on *q*^*th*^ order absolute moment of a time series independent of *q*. Estimating 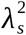 based on 0.2^th^ moment (*q*= 0.2) emphasizes the center part of the PDF, reducing the effects of extreme deviations due to heavy-tails and kurtosis. We used 0.2^th^ moment because estimates of for 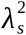 a time series of ∼ 6000 samples are more accurate at lower values of *q* [79].

**Figure 2.**
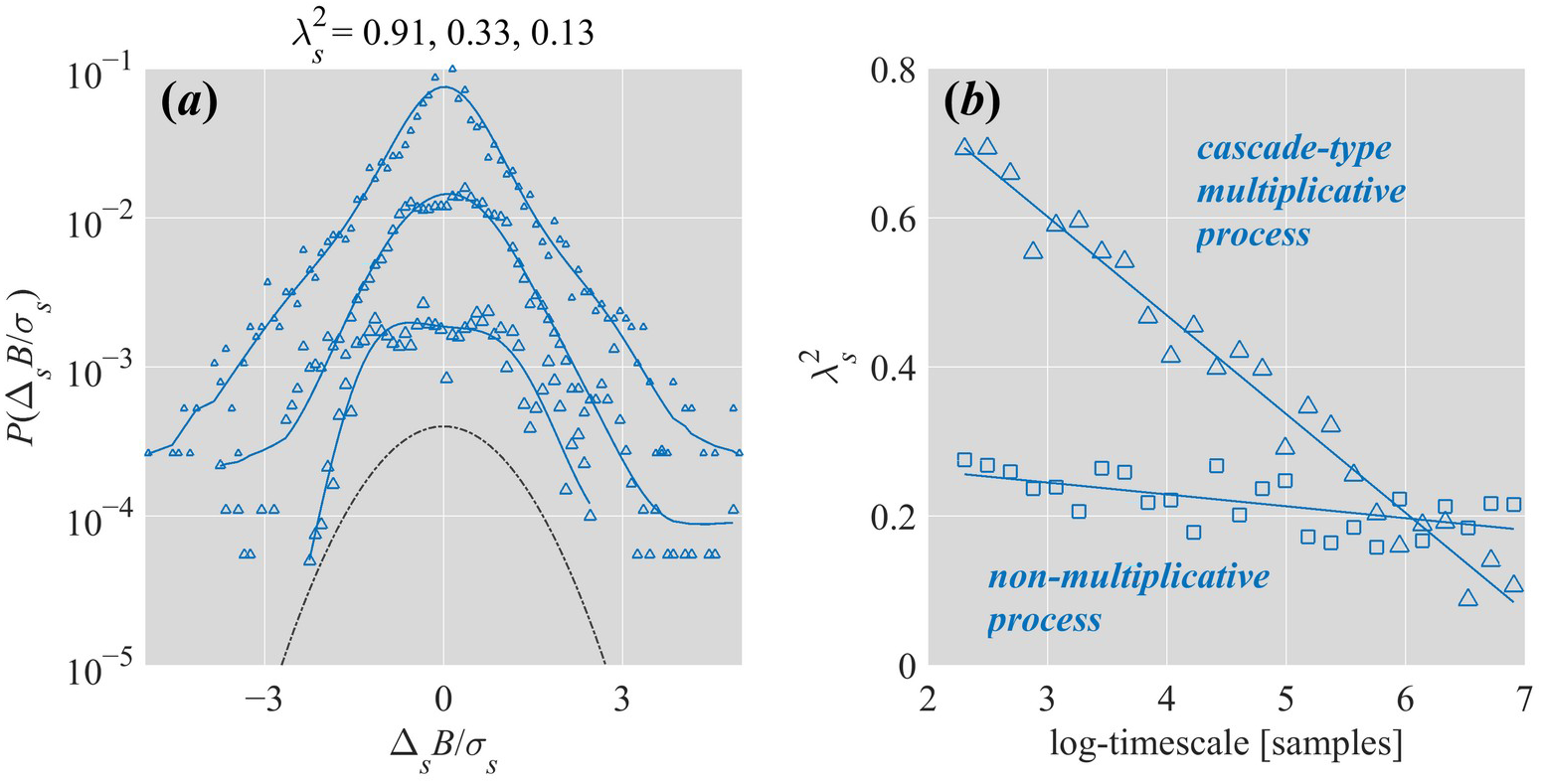
Schematic illustration of non-Gaussianity index 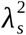. (*a*) The relationship between 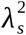 and shapes of PDF plotted in linear-log coordinates for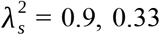, and 0.13 (from top to bottom). For clarity, the PDFs have been shifted vertically. As 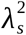 increases, the PDF becomes increasingly peaked and fat-tailed. As *λs* decreases, the PDF increasingly resembles the Gaussian (dashed line), assuming a perfect Gaussian as 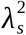 approaches 0. (*b*) In a cascade-type multiplicative process yield the inverse relationship 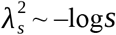.

Cascade-type multiplicative process yield the inverse relationship 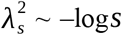 (figure 2*b*) [55]. For the present purposes, we quantified 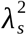 for each original CoP PED series and corresponding IAAFT surrogate at timescales 5 to 1000 samples (i.e., 50 ms to 10 s) at steps of 5 samples (50 ms).

To test explicitly for lognormality in postural fluctuations, we used Akaike Information Criterion (AIC) weights obtained via the more common maximum likelihood estimation (MLE) to determine whether the PDF at each timescale was a power law, lognormal, exponential, or gamma distribution. We include this analysis also to complement multiscale PDF analysis to portray the analytical effect of sensitivity to different portions of the distribution.

### 2.6. Statistical analysis

To test for crossovers in non-Gaussianity, a linear mixed-effect (LME) model using lmer() function in package *lme4* [80] for *R* [81] tested 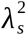 vs. log-timescale curves for orthogonal linear and quadratic polynomials, for interactions with grouping variables (Surface × Vision × Age-group × Original, where Original encoded differences in original series from surrogates) and with indices of postural control (Section 2.4). Statistical significance was assumed at the alpha level of 0.05 using package *lmerTest* [82]. To test how lognormality changed with log-timescale, a generalized linear mixed-effect (GLME) fit changes in Lognormality as a dichotomous variable using orthogonal linear, quadratic, and cubic polynomials and tested interaction effects of grouping variables (Surface × Vision × Age × Original) with those polynomials using glmer() function in package *lme4* [80].

## 3. Results

Figures 3 and 4 provide a schematic of multiscale PDF characterization of postural sway in representative young and old participants standing quietly for 60 s on stable and unstable surfaces with eyes closed (figure 3) and eyes open (figure 4).

**Figure 3.**
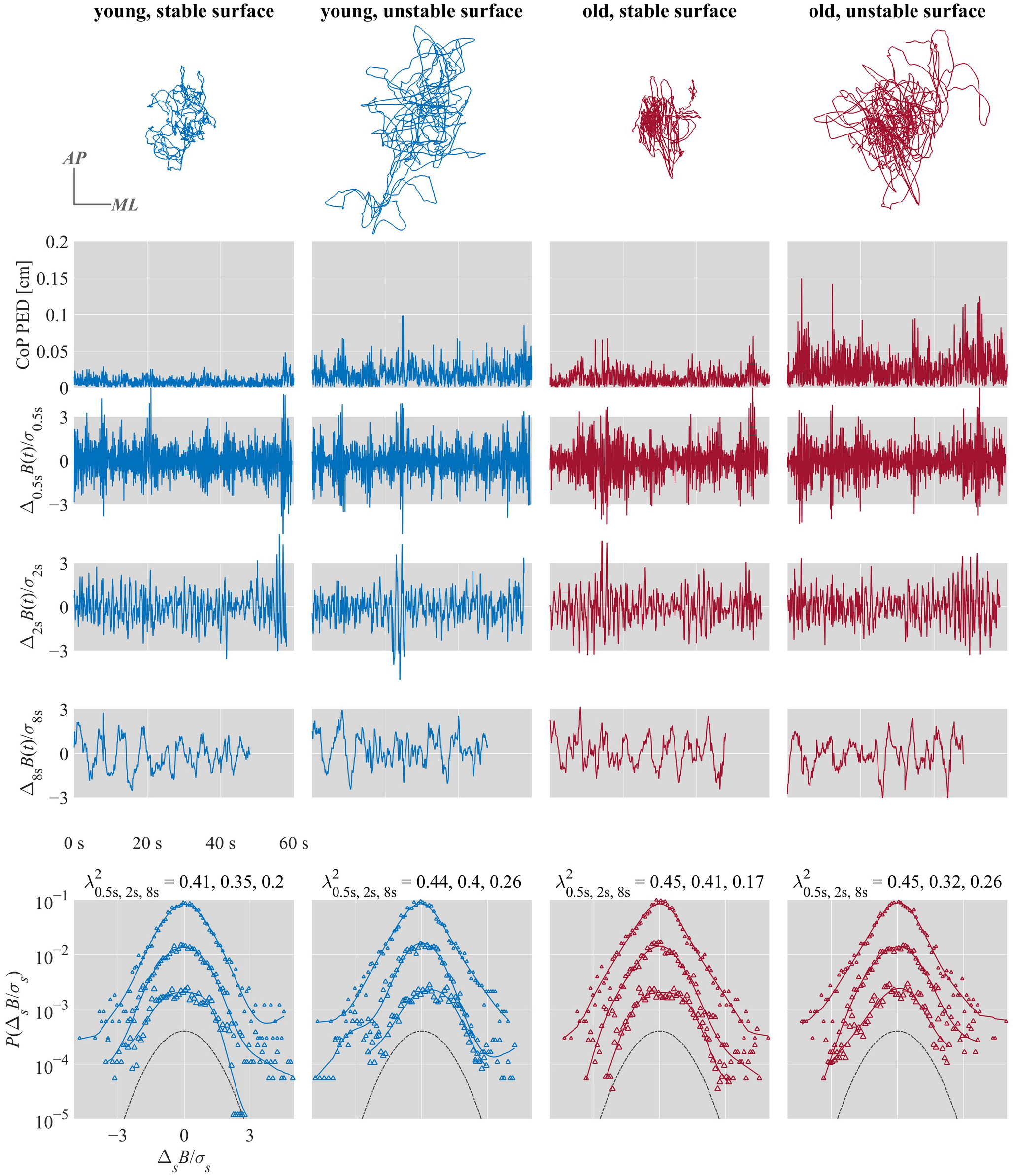
Multiscale PDF characterization of postural sway in representative young and old participants standing quietly for 60 s on stable and unstable surfaces with eyes open. From top to bottom: CoP trajectories along the anterior-posterior (*AP*) and medial-lateral (*ML*) axes. CoP PED series. {*Δ*_*s*_ *B* (*i*)} for *s* = 0.5, 2, and 8 s. Standardized PDFs (in logarithmic scale) of {*Δ*_*s*_ *B* (*i*)} for *s* = 0.5, 2, and 8 s (from top to bottom), where *σ* _*s*_ denotes the *SD* of {*Δ*_*s*_ *B* (*i*)}.

**Figure 4.**
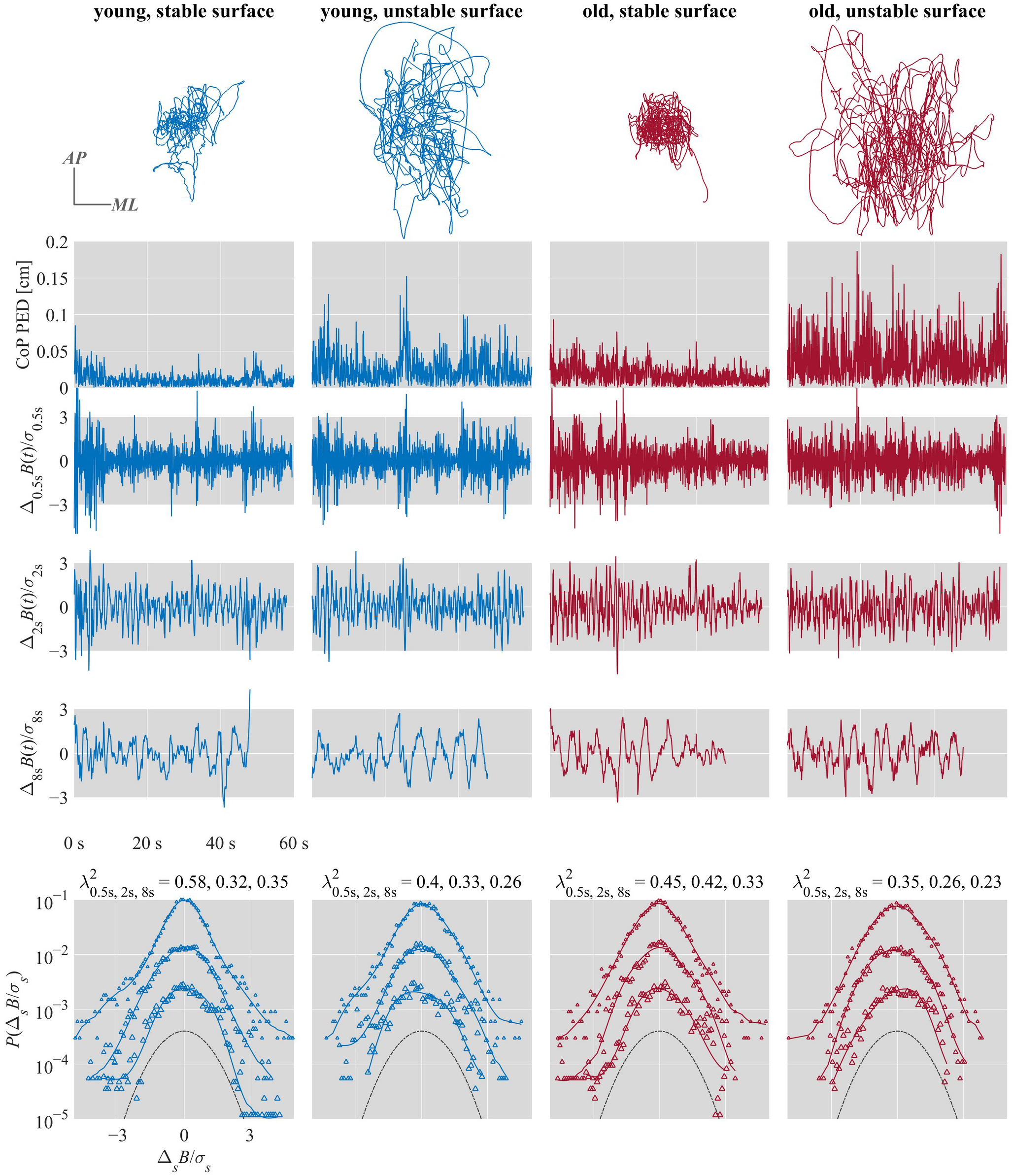
Multiscale PDF characterization of postural sway in representative young and old participants standing quietly for 60 s on stable and unstable surfaces with eyes closed. From top to bottom: CoP trajectories along the anterior-posterior (*AP*) and medial-lateral (*ML*) axes. CoP PED series. {*Δ*_*s*_ *B* (*i*)} for *s* = 0.5, 2, and 8 s. Standardized PDFs (in logarithmic scale) of {*Δ*_*s*_ *B* (*i*)} for *s* = 0.5, 2, and 8 s (from top to bottom), where *σ* _*s*_ denotes the *SD* of {*Δ*_*s*_ *B* (*i*)}.

### 3.1. Hypothesis-1

#### 3.1.1. 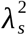 -vs.-log-timescale curves for stable postural conditions

Surface stability elicited nonlinearity in non-Gaussianity with timescale, that is, low short-scale 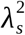, increasing up to a maximum roughly at 75 samples (750 ms), and decreasing with subsequently longer timescales (table S1; figure 5a). 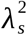 showed stronger linear decrease (*b* = –13.37, *p* < 0.001) and downward-facing parabolic shape (*b* = –2.57, *p* < 0.001). This profile of 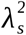-vs.-log-timescale curves held for original series and not for surrogates without original nonlinear interactions across timescales.

**Figure 5.**
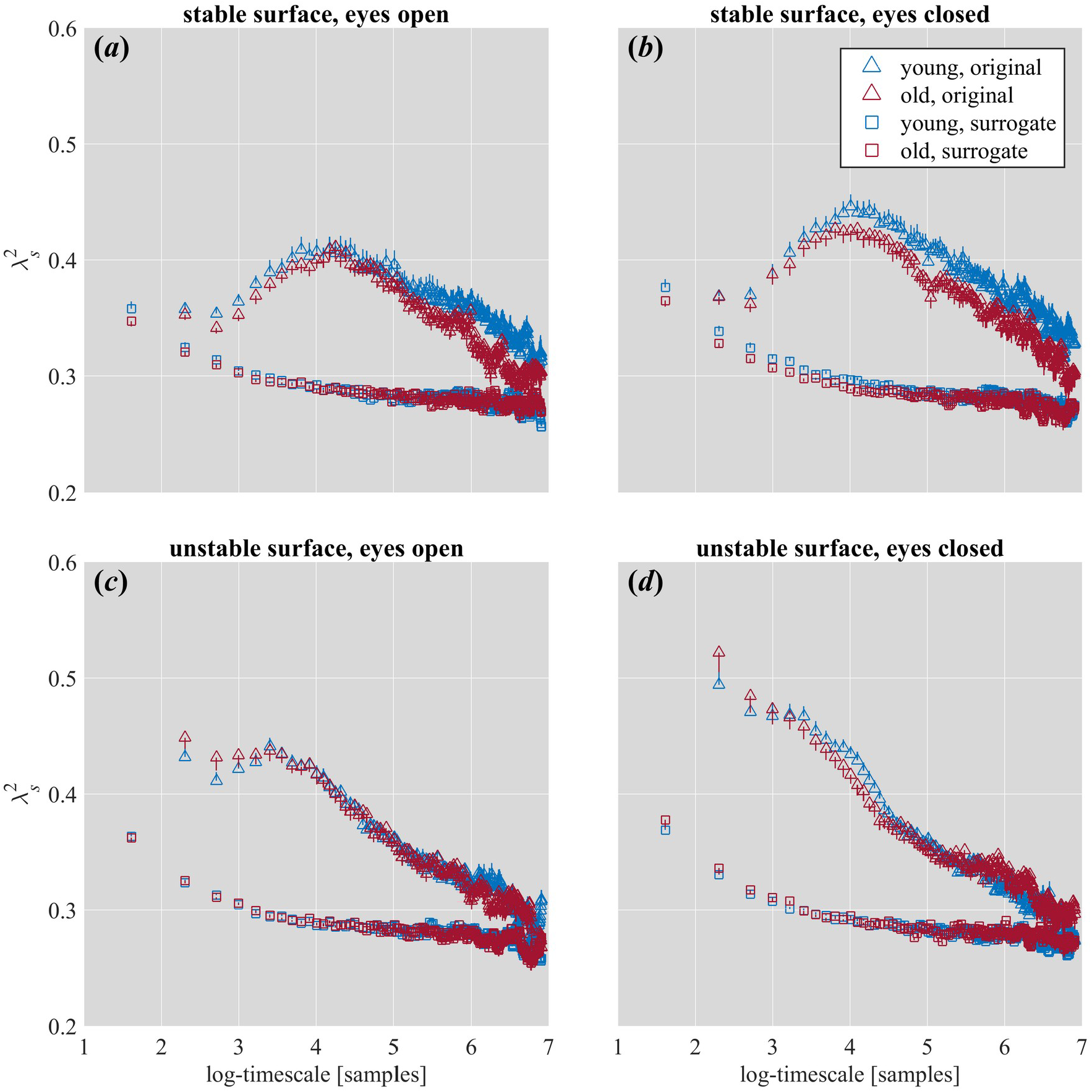
Log-timescale dependence of the non-Gaussianity index 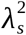 for the young and old participants standing quietly for 60 s. (*a*) Stable surface, eyes open. (*b*) Stable surface, eyes closed. (*c*) Unstable surface, eyes open. (*d*) Unstable surface, eyes closed. Vertical bars indicate ±1*s*.*e*.*m*. of the group averages (young, *n* = 27; old, *n* = 22).

#### 3.1.2. The growth of lognormality in postural sway with log-timescale for stable postural conditions

For the most stable postural conditions (i.e., stable surface, eyes open), heaviness of PDF tails aligned with results from multiscale PDF analysis of non-Gaussianity (table S2; figure 6 *a*). The original CoP PED series exhibited less lognormality overall than the respective surrogates (*b* = –1.75, *p* < 0.001). The stable-surface and eyes-open conditions yielded a cubic-polynomial increase in lognormality across log-timescale. Specifically, the CoP PED series showed positive linear (*b* = 296.15, *p* < 0.001) and quadratic (*b* = 161.98, *p* < 0.001) but negative cubic (*b* = –209.16, *p* = < 0.001) growth with log-timescale. However, a striking result is notably heavier PDF tails (i.e., more likely to be lognormal) for surrogates without original nonlinear interactions across timescales.

**Figure 6.**
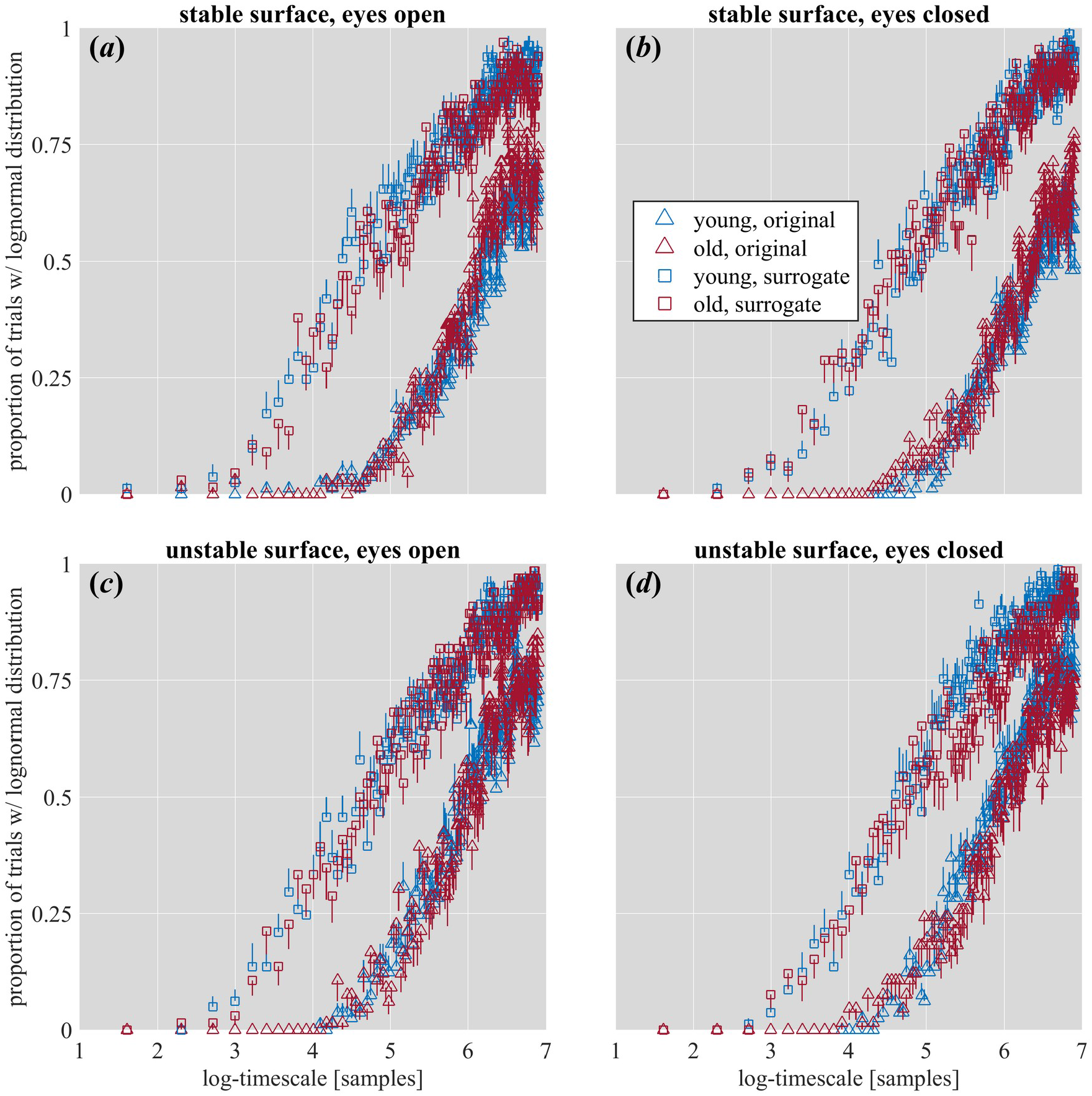
Log-timescale dependence of the mean proportion of trials with a lognormal distribution for the young and old participants standing quietly for 60 s. (*a*) Stable surface, eyes open. (*b*) Stable surface, eyes closed. (*c*) Unstable surface, eyes open. (*d*) Unstable surface, eyes closed. Vertical bars indicate ±1*s*.*e*.*m*. of the group averages (young, *n* = 27; old, *n* = 22).

### 3.2. Hypothesis-2

#### 3.2.1. 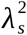-vs.-log-timescale curves for less stable postural conditions

Less stable postural conditions consistently showed increased non-Gaussianity and more cascade-like linear decay of with log-timescale in original series as compared to linear surrogates (table S1; figures 5 *b* and 5*d*). First, young participants showed a significantly weaker linear decrease in stable-surface with longer timescales (*b* = 3.74, *p* < 0.001), indicating that stronger linear descent of 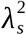 is attributable to weaker stability with age. However, changes in the nonlinearity of 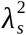-vs.-log-timescale curves did not depend on age alone or differ between the eyes-open and eyes-closed conditions (*p* > 0.05).

Standing on the unstable surface with eyes open showed significantly less of the downward-facing parabolic shape (figure 5*a*) as when standing on the stable surface with eyes open (*b* = 8.64, *p* < 0.001; figure 5*c*).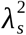 showed stronger linear decay with log-timescale for the older participants with eyes open (*b* = –5.37, *p* < 0.001). Combining surface instability with eyes closed elicited characteristic evidence of scale-invariant cascades, that is, stronger linear decreases (*b* = –1.24, *p* < 0.001) and weaker quadratic forms (*b* = 3.26, *p* < 0.001) of 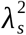 -vs.-log-timescale irrespective of age (figure 5*d*). Young participants exhibited even stronger linear decay (*b* = –2.26, *p* = 0.001) but slightly less reversal of the downward-facing parabolic decay in 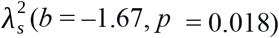. Young participants also showed an overall increase in 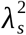 with eyes closed (*b* = 0.0069, *p* < 0.001).

#### 3.2.2. The growth of lognormality in postural sway with log-timescale for less stable postural conditions

The relationship between feedback-based control and PDF tails began to unravel, as results about non-Gaussianity and lognormality of postural fluctuations began to diverge, although some changes in lognormality aligned with the results about non-Gaussianity. For instance, young participants showed less lognormality overall (*b* = –0.42, *b* < 0.001; figure 6), indicating that greater lognormality is attributable to weaker stability with age. However, factors that should have destabilized posture did not increase lognormality—for example, closing eyes also reduced the likelihood of lognormality (*b* = –0.41, *p* < 0.001; figures 6*b* and 6*d*), signifying a departure of PDF tail heaviness from non-Gaussian evidence of cascade strength.

Surface instability and its combination with closed eyes and age produced progressively stronger signatures of cascade structure, but the effects of these factors on PDF tail heaviness were much smaller and mixed in direction. Surface instability increased lognormality for young participants to a greater degree than for old participants (*b* = 0.37, *p* = 0.002; figures 6*c* and 6*d*) as well as increased lognormality even more for all participants with eyes closed (*b* = 0.62, *p* < 0.001). Surface instability weakened quadratic growth with eyes open (*b* = –248.63, *p* = 0.042; figure 6c) but strengthened it with eyes closed (*b* = 417.63, *p* = 0.041; figure 6d). Effects of surface instability depended on age, with closed eyes accentuating the cubic form for the old participants (*b* = –231.37, *p* = 0.027) but reversing it for young participants (*b* = 353.95, *p* = 0.040).

### 3.3. Hypothesis-3: Effects of indices of endogenous postural control on 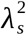-vs.-log-timescale curves

All linear and nonlinear indices of postural control showed predictive effects on 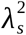 (table S1). 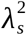Typically decreased with increases in CoP_*Mean* (*b* = –41.67, *p* < 0.001), CoP_*SD* (*b* = 31.01, *p* < 0.001), CoP_*SampEn* (*b* = –0.01, *p* < 0.001), and CoP_*MSE* (*b* = –0.08, *p* < 0.001), and increased with increases in CoP_*RMSE* (*b* = 51.76, *p* < 0.001). 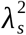 also increased with both the strength of temporal correlations or fractality (i.e., CoP_ *H*_fGn_; *b* = 0.08, *p* < 0.00) and more variation in these temporal correlations or multifractality (i.e., CoP_dAlpha; *b* = – 0.05, *p* < 0.00). Nonetheless, in the tradeoff between temporal correlations, greater ‘original-minus-surrogate’ differences in both fractality and multifractality diminished (CoP_*H*_fGn__diff_OS: *b* = 2.72, *p* < 0.001; CoP_dAlpha_diff_OS: *b* = –2.72, *p* < 0.001). This result confirms the fact that evidence of multifractality with a relatively weak difference from the surrogates suggests heavy-tailed distributions and thus non-Gaussianity [59,60]. However, this model, including only the main effects of these indices, speaks only to average differences in intercept and not to the trajectory of across log-timescale.

Including the interactions of all indices of postural sway with linear and quadratic terms in the prior model significantly improved the model fit (*χ*_2423_ = 32699, *p* < 0.001; table S3). Table S3 details the full model, but in short, it illustrated two major points: first, most of the variability in the quadratic form of the 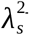 vs.-log-timescale curves depended on these indices. This newer model with interactions of these indices with log-timescale had 69 significant quadratic effects (*p* < 0.05), and all but one significant quadratic effects were interactions of these indices with log-timescale. That is, the parabolic tempering of non-Gaussianity across log-timescale in stable-surface condition was mostly a function of endogenous postural control.

## 4. Discussion

Multiscale PDF analysis was used to test whether unperturbed postural sway exhibits weaker and stronger evidence of cascade dynamics at relatively short and relatively long timescales. Our major finding is that of a crossover in non-Gaussianity profiles of postural sway with log-timescale. This crossover likely reflects a transition between distinct control processes enlisting feedback on short timescales but not long timescales. The present work joins a growing enterprise in attempting to characterize diverse control processes through nonlinear methods. Perturbing postural sway with unstable support surface or with closing eyes reveals the capacity of non-Gaussian cascade dynamics to spread from the longer timescales into shorter timescales in both old and young participants.

The present findings offer new grounds for doubting whether the traditional open-/closed-loop interpretation of control is fundamental [1,2,83,84]. Sensory processes and postural corrections do not necessarily observe a firm divide between fast and slow processes [85]. Instead, they reflect contingent scale-dependencies of postural corrections emerging from constraints subject to task manipulation. Task manipulation specifically working to destabilize posture—whether by standing on an unstable surface or closing eyes—reveal rich cascade-like dynamics in postural control [50–52]. Cascade dynamics afford richly interactive capacity for movement organization to emerge through the coordination in accordance with constraints at multiple scales [86,87].

Non-Gaussianity complements the existing understanding that healthy human postural sway exhibits complexity quantifiable through various means (entropy [69], fractality [88], and multifractality [78]). The present work shows that the cascade structure of healthy human postural sway varies not simply with the extrinsic manipulation of task constraints but also with the postural system’s endogenous complexity. Indices of postural sway governed the appearance or disappearance of crossovers, suggesting that tempering of non-Gaussianity across log-timescale in stable condition is mostly a function of endogenous postural control. This pliability of a crossover between inertial and feedback-based control could have life-and-death consequences for healthy functioning (e.g., preventing falls). The present work suggests that an important direction for future research would be using multiscale PDF as a lens on the healthy maintenance of upright standing, particularly as a balance between endogenous control and exogenous constraints.

## Supplementary materials

Supplementary Table S1. Coefficients of linear mixed-effect (LME) model examining the effects of Surface × Vision × Age-group on 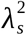 vs. log-timescale curves.

Supplementary Table S2. Coefficients of generalized linear mixed-effect (GLME) model examining the effects of Surface × Vision × Age-group × Type on the growth of lognormality in postural sway with log-timescale.

Supplementary Table S3. Coefficients of LME model examining the effects of Surface × Vision × Age-group on 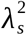 vs. log-timescale curves for orthogonal linear and quadratic polynomials, for interactions of grouping variables (Surface × Vision × Age-group × Type) and with indices of postural control.

## Author contributions

M.M. and D.G.K-S. conceived and designed research; M.M. and D.G.K-S. analyzed data; M.M. and D.G.K-S. interpreted results of experiments; M.M. prepared figures; M.M. and D.G.K-S. drafted manuscript; M.M. and D.G.K-S. edited and revised manuscript; M.M. and D.G.K-S. approved final version of manuscript.

